# Trypsin treatment unlocks barrier for zoonotic coronaviruses infection

**DOI:** 10.1101/768663

**Authors:** Vineet D. Menachery, Kenneth H. Dinnon, Boyd L. Yount, Eileen T. McAnarney, Lisa E. Gralinski, Andrew Hale, Rachel L. Graham, Trevor Scobey, Simon J. Anthony, Lingshu Wang, Barney Graham, Scott H. Randell, W. Ian Lipkin, Ralph S. Baric

## Abstract

Traditionally, the emergence of coronaviruses (CoVs) has been attributed to a gain in receptor binding in a new host. Our previous work with SARS-like viruses argued that bats already harbor CoVs with the ability to infect humans without adaptation. These results suggested that additional barriers limit the emergence of zoonotic CoV. In this work, we describe overcoming host restriction of two MERS-like bat CoVs using exogenous protease treatment. We found that the spike protein of PDF2180-CoV, a MERS-like virus found in a Ugandan bat, could mediate infection of Vero and human cells in the presence of exogenous trypsin. We subsequently show that the bat virus spike can mediate infection of human gut cells, but is unable to infect human lung cells. Using receptor-blocking antibodies, we show that infection with the PDF2180 spike does not require MERS-CoV receptor DPP4 and antibodies developed against the MERS spike receptor-binding domain and S2 portion are ineffective in neutralizing the PDF2180 chimera. Finally, we found that addition of exogenous trypsin also rescues replication of HKU5-CoV, a second MERS-like group 2c CoV. Together, these results indicate that proteolytic cleavage of the spike, not receptor binding, is the primary infection barrier for these two group 2c CoVs. Coupled with receptor binding, proteolytic activation offers a new parameter to evaluate emergence potential of CoVs and offer a means to recover previously unrecoverable zoonotic CoV strains.

**Importance:** Overall, our studies demonstrate that proteolytic cleavage is the primary barrier to infection for a subset of zoonotic coronaviruses. Moving forward, the results argue that both receptor binding and proteolytic cleavage of the spike are critical factors that must be considered for evaluating the emergence potential and risk posed by zoonotic coronaviruses. In addition, the findings also offer a novel means to recover previously uncultivable zoonotic coronavirus strains and argue that other tissues, including the digestive tract, could be a site for future coronavirus emergence events in humans.

## Introduction

Since the beginning of the 21^st^ century, public health infrastructures have been required to periodically respond to new and reemerging zoonotic viral diseases, including influenza, Ebola, and Zika virus outbreaks (1). Severe acute respiratory syndrome coronavirus (SARS-CoV), the first major outbreak of the century, highlighted the global impact of a newly emerging virus in the context of expanding development, increased globalization, and poor public health infrastructures (2-4). A decade later, the emergence and continued outbreaks of the Middle East respiratory syndrome coronavirus (MERS-CoV) further illustrate the ongoing threat posed by circulating zoonotic viruses (5). Together, the outbreaks of the early part of this century argue that continued preparations and vigilance are needed to maintain global public health.

Despite their spontaneous emergence, several research approaches to rapidly respond and even predict outbreak strains already exist. During the MERS-CoV outbreak, our group and others were able to leverage reagents generated against related group 2C coronaviruses, HKU4- and HKU5-CoV (6, 7). These reagents, created independent of viable virus replication, provided valuable insights and models for testing serologic responses during the early stages of the MERS-CoV outbreak. Similarly, reverse genetics systems permitted the exploration of zoonotic coronaviruses (8); using the known SARS spike/ACE2 receptor interaction, chimeric viruses containing the backbones of bat CoVs were generated to evaluate the efficacy of both vaccines and therapeutics (9-12). The inverse approach placed the zoonotic spike proteins in the context of the epidemic SARS-CoV backbone (13, 14). These studies provided insight into potential threats circulating in bats as well as the efficacy of current therapeutic treatments (15). While far from comprehensive, the results indicated that these approaches, reagents, and predictions may prove useful in preparations for future CoV outbreaks.

In this study, we extend examination of zoonotic viruses to a novel MERS-like CoV strain isolated from a Ugandan bat, PDF-2180 CoV (MERS-Uganda). Our initial attempt to cultivate a chimeric MERS-CoV containing the Ugandan MERS-like spike produced viral sub-genomic transcripts, but failed to result in infectious virus after electroporation (16). However, in the current study, we demonstrate that exogenous trypsin treatment produced high-titer virus capable of plaque formation and continued replication. The chimeric Ugandan MERS-like spike virus could replicate efficiently in both Vero and Huh7 cells in the context of trypsin-containing media, but failed to produce infection of either continuous or primary human respiratory cell cultures. Importantly, the MERS-Uganda chimeric virus successfully infected cells of the human digestive tract, potentially identifying another route for cross-species transmission and emergence. Notably, blockade of human DPP4, the receptor for MERS-CoV, had no significant impact on replication of the chimeric MERS-Uganda virus, suggesting the use of an alternative receptor. Similarly, addition of trypsin also rescued replication of full-length HKU5-CoV, a related group 2C bat CoV, and showed no replication defect during DPP4 blockade. Together, the results indicate that proteolytic activation of the spike protein is a potent constraint to infection for zoonotic CoVs and expands the correlates for CoV emergence beyond receptor binding alone.

## Results

Utilizing the MERS-CoV infectious clone (17), we previously attempted to evaluate the potential of the PDF-2180 CoV to emerge from zoonotic populations. Replacing the wild-type MERS-CoV spike with the PDF-2180 spike produced a virus capable of generating viral transcripts in vitro, but not sustained replication (16). These results suggested that the significant amino acid differences observed within the receptor-binding domain precluded infection of Vero cells. However, amino acid changes were not confined only to the receptor-binding domain (RBD); highlighting changes between the Uganda spike on the MERS-CoV trimer revealed significant differences throughout the S1 region of spike (**Fig. 1A & B**). While the S2 remained highly conserved (**Fig. 1C**), changes in the C- and N-terminal domains of S1, in addition to the RBD, may also influence entry and infection compatibility. Notably, recent reports had also indicated differential protease cleavage of wild-type MERS-CoV based on cell types, suggesting that spike processing influences docking and entry of pseudotyped virus (18). To explore if spike cleavage impaired infectivity, we evaluated MERS-Uganda virus replication in the presence of trypsin-containing media. The addition of trypsin to the chimeric virus resulted in cytopathic effect, fusion of the Vero monolayer, formation of plaques under a trypsin-containing overlay, and collection of high-titer infectious virus stock (**Fig. 1D**). The requirement for trypsin complicated these studies due to cell toxicity; to overcome this issue, we utilized both trypsin-adapted Vero cells and a MERS-Uganda chimera encoding RFP in place of ORF5, similar to a previously generated MERS-CoV reporter virus (17). Following MERS-Uganda infection, cultures with trypsin containing media showed evidence for replication of viral genomic RNA (**Fig. 1E**). Similarly, the nucleocapsid protein was only observed in the presence of exogenous trypsin following infection with the MERS-Uganda chimera (**Fig. 1F**). Notably, wild-type MERS-CoV was also augmented in the presence of trypsin with increased genomic RNA and nucleocapsid protein relative to no trypsin control (**Fig. 1E &F**). Examination of RFP signal confirmed these RNA and protein results (**Fig. 1G**), as RFP was only observed in MERS-Uganda chimeric infection in the presence of trypsin. Similarly, RFP expression was more robust in trypsin-treated cells following MERS-CoV infection. Together, these data indicate that the PDF-2180 spike can mediate infection of Vero cells in a trypsin-dependent manner.

**Figure 1.**
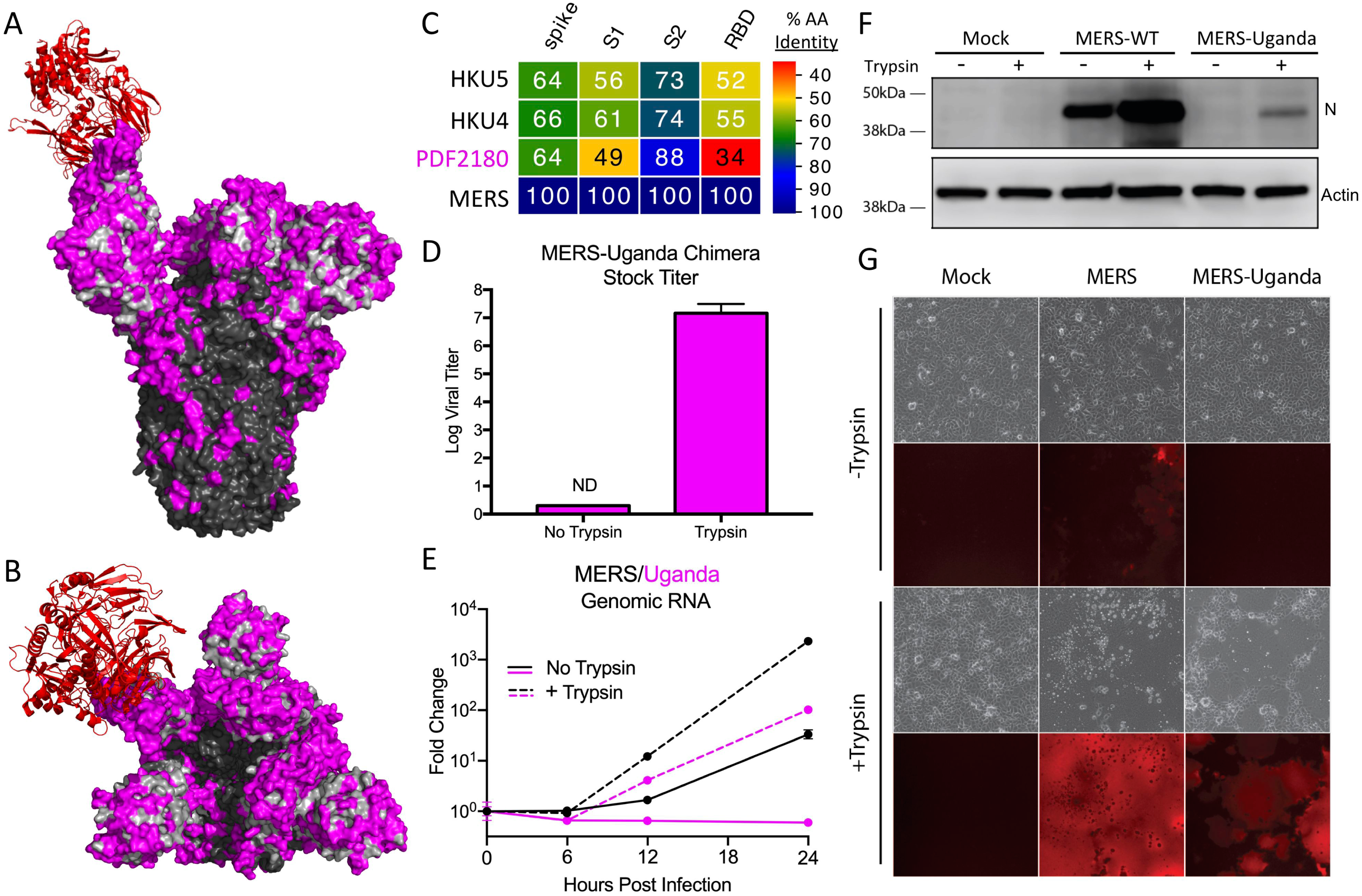
Exogenous trypsin rescues MERS-Uganda spike replication. A & B) Structure of the MERS-CoV spike trimer in complex with the receptor human DPP4 (red) from the A) side and B) top. Consensus amino acids are outlined for the S1 (grey) and S2 (black) domains, with PDF-2180 differences noted in magenta. C) Spike protein sequences of the indicated viruses were aligned according to the bounds of total spike, S1, S2, and receptor-binding domain (RBD). Sequence identities were extracted from the alignments, and a heatmap of sequence identity was constructed using EvolView (http://www.evolgenius.info/evolview) with MERS-CoV as the reference sequence. D) MERS-Uganda chimera stocks were grown in the presence or absence of trypsin and were quantitated by plaque assay with a trypsin-containing overlay (n = 2). E) Expression (qRT-PCR) of MERS-CoV (black) and MERS-Uganda (magenta) genomic RNA following infection of Vero cells in the presence or absence of trypsin (n=3 for each time point). F) Protein expression of MERS-CoV nucleocapsid (N) and actin 18 hours post-infection of Vero in the presence or absence of trypsin in the media. G) Phase-contrast and RFP expression microscopy in Vero cells infected with MERS-CoV, MERS-Uganda spike chimera, or mock in the presence or absence of trypsin.

### MERS-Uganda spike replicates in human cells

Having demonstrated infection and replication, we next sought to determine the capacity of MERS-Uganda chimeric virus to grow in human cells. Previously, MERS-CoV had been shown to replicate efficiently in Huh7 cells (19). Using the Huh7 liver cell line, infection with MERS-Uganda RFP chimeric virus resulted in RFP-positive cells and cell fusion (**Fig. 2A**). In contrast, while a few RFP-positive cells were observed in the non-trypsin-treated group, neither expanding RFP expression nor cytopathic effect were seen in the absence of trypsin. Our observation may have been the result of residual trypsin activity from the undiluted virus stock, resulting in low-level infection. Exploring further, N protein analysis by Western blot indicated that the PDF-2180 spike chimera could produce significant viral proteins in the presence of trypsin (**Fig. 2B**); only low levels of protein were observed in the control-treated infection. While replication of the MERS-Uganda chimera was not equivalent to that of wild-type MERS-CoV, the results clearly demonstrate the capacity of the PDF-2180 spike to mediate infection of human cells in the presence of trypsin.

**Figure 2.**
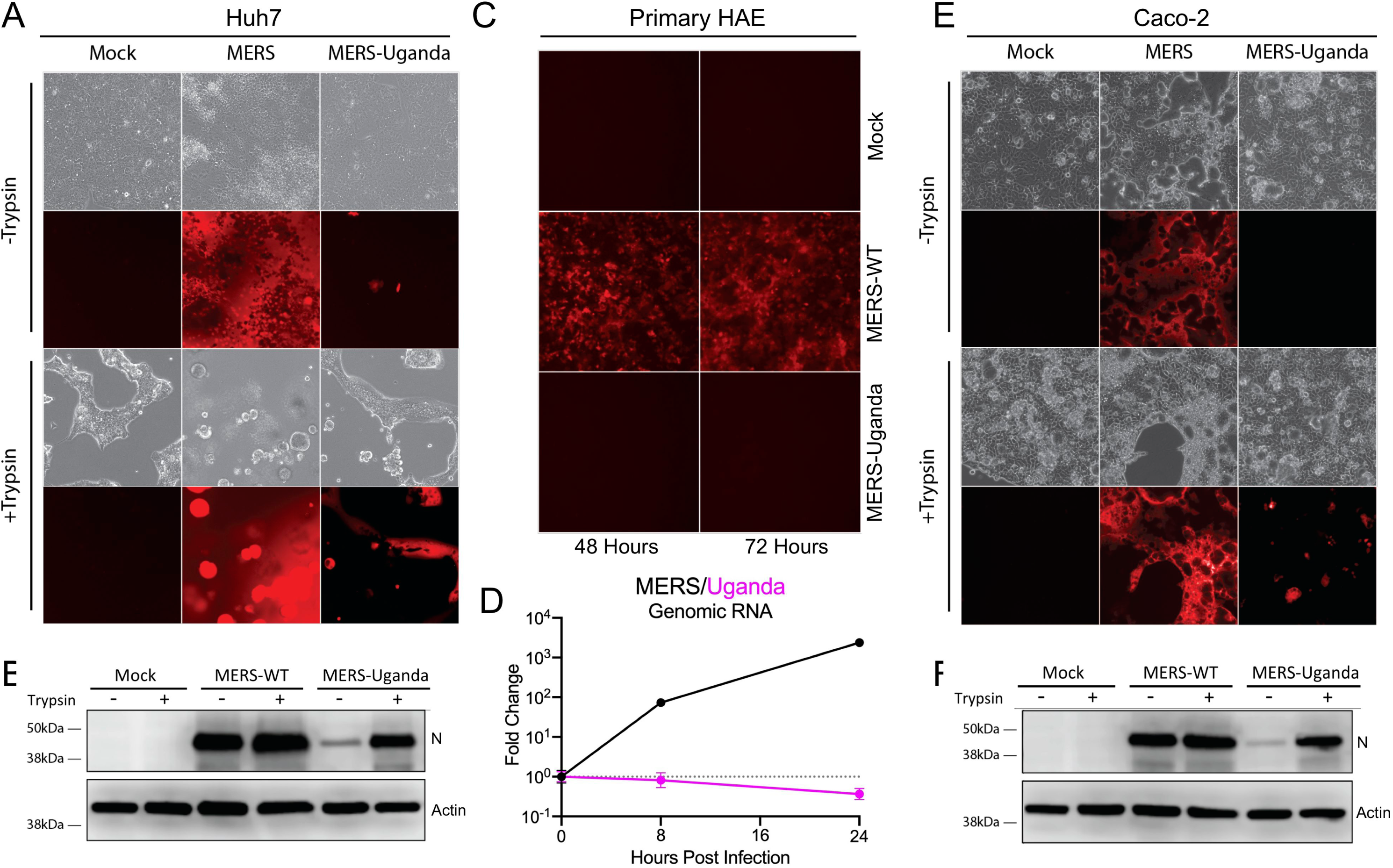
MERS-Uganda spike chimera replicates in human cells. A & B) Huh7 cells were infected with MERS-CoV or MERS-Uganda chimeric viruses, showing A) microscopy images of cell monolayer and RFP expression with and without trypsin treatment and B) N protein expression following infection of Huh7 cells in the presence or absence of trypsin. C & D) Primary HAE cultures were infected with MERS-CoV or MERS-Uganda chimera, showing C) RFP expression and D) genomic viral RNA following infection (n = 3 for 8, 24 HPI). E & F) Caco-2 cells were infected with MERS-CoV or MERS-Uganda chimeric viruses expressing RFP, showing E) microscopy images of cell monolayer and RFP expression with and without trypsin treatment and F) N protein expression following infection of Caco-2 cells in the presence or absence of trypsin.

We next examined the capacity of MERS-Uganda spike to infect human respiratory cells, the primary targets of SARS-CoV, MERS-CoV, and other common-cold human CoVs. Using Calu3 cells, a human lung epithelial cell line, we observed robust replication of wild-type MERS-CoV based on RFP expression, consistent with previous studies (17). However, no evidence of infection was noted in MERS-Uganda-infected Calu3 cells in the presence or absence of trypsin. We subsequently explored primary human airway epithelial (HAE) cultures. Grown on an air–liquid interface, HAE cultures have a propensity to facilitate improved infections of several human CoVs and may be more permissive for infection with the PDF-2180 spike chimera (20). However, infection with PDF-2180 spike-containing virus showed no evidence of RFP expression, even after several trypsin washes of the apical surface (**Fig. 2C**). Similarly, RNA expression analysis found no evidence for accumulation of viral genomic RNA, indicating no evidence for replication in HAE cultures (**Fig. 2D**). In contrast, wild-type MERS-CoV efficiently infects these HAE cultures, as demonstrated by both RFP expression and viral genomic RNA accumulation. Together, the Calu3 and HAE results suggest that the PDF-2180 spike is unable to infect respiratory cells in humans, even in the presence of exogenous trypsin.

We next evaluated the capacity of the PDF-2180 chimera to infect cells of the digestive tract. While uncommon in humans, several animal CoVs have been shown to cause severe disease via the enteric pathway (21, 22). In addition, most bat CoV sequences, including PDF-2180-CoV, were isolated from bat guano samples, suggesting an enteric etiology. Importantly, the presence of trypsin and other soluble host proteases in the digestive tract may facilitate infection with PDF-2180 spike in humans. To test this question, we infected Caco-2 cells, a human epithelial colorectal adenocarcinoma cell line, with wild-type MERS-CoV and MERS-Uganda spike chimera in the presence or absence of trypsin (**Fig. 2E**). For MERS-CoV, infection of Caco-2 cells resulted in robust infection and spread with or without trypsin in the media. For the MERS-Uganda chimera, the addition of trypsin facilitated infection with abundant RFP-positive Caco-2 cells; however, infection was not as robust as in the wild-type MERS-CoV infection. Examination of N protein by Western blot indicated that the MERS-Uganda spike could produce infection in Caco-2 cells, but confirmed replication at levels lower than with wild-type MERS-CoV (**Fig. 2F**). Together, the results indicate that human cells, including gut cells, can support infection with MERS-Uganda chimera in the presence of trypsin.

### MERS-Uganda spike does not use DPP4 for entry

The absence of infection of human respiratory cells coupled with significant changes in the RBD suggested that MERS-Uganda does not utilize the MERS-CoV receptor, human DPP4, for entry (16). To explore this question, we utilized antibodies to block DPP4 in Vero cells to determine the effect on MERS-Uganda chimeric virus replication. As expected, anti-DPP4 antibody successfully ablated replication of wild-type MERS-CoV in both the presence and the absence of trypsin treatment, as measured by both RFP and N protein expression (**Fig. 3A & B**). In contrast, the human DPP4-blocking antibody had no impact on infection with the MERS-Uganda chimera virus in the presence of trypsin, confirming that the MERS-CoV receptor is not required to mediate infection with the PDF-2180 spike. Together, these results indicate that while the MERS-Uganda spike infects human cells, it does not require human DPP4 to mediate infection.

**Figure 3.**
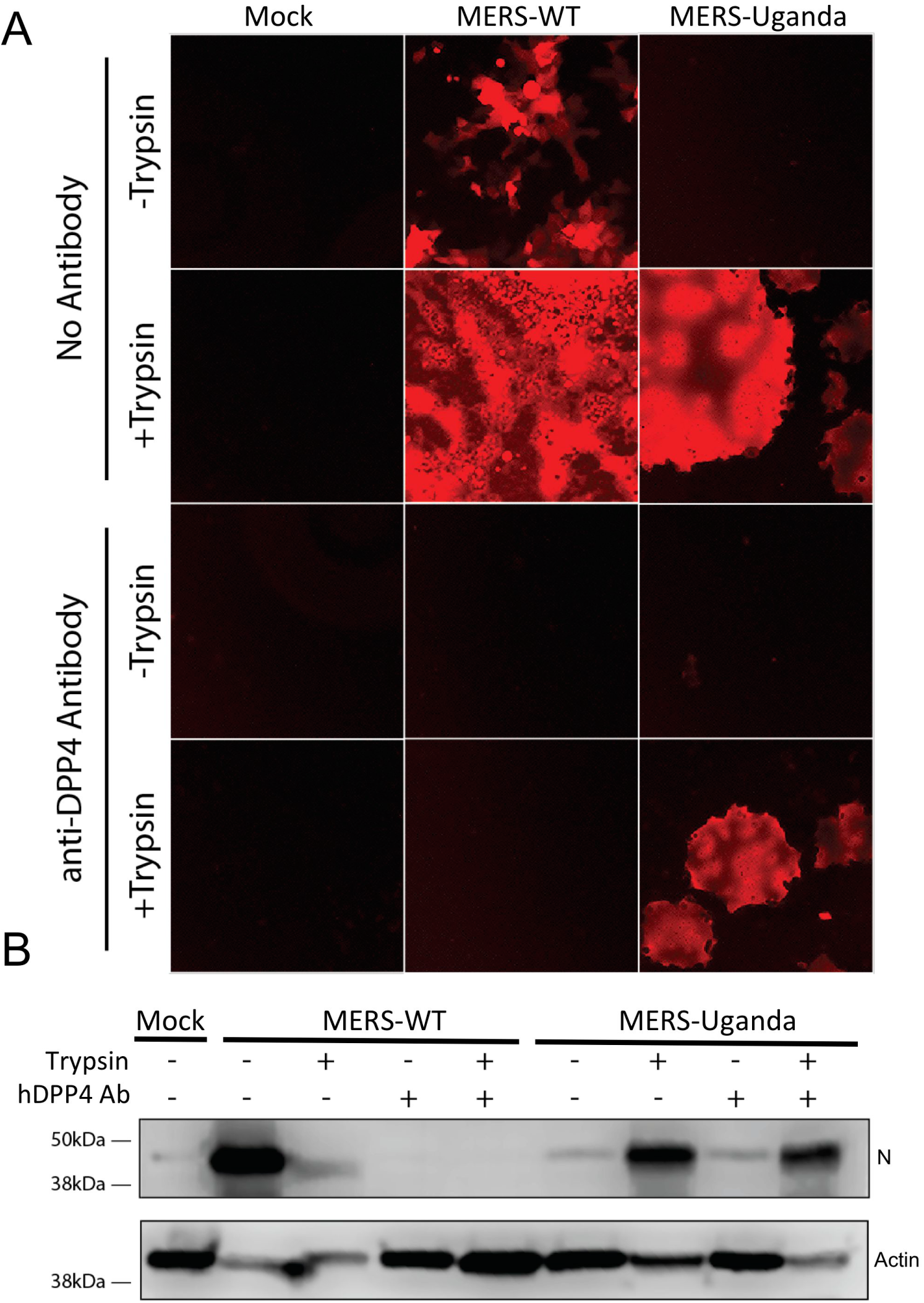
MERS-Uganda spike does not utilize DPP4 for infectfsion. A & B) Vero cells were infected with MERS-CoV or MERS-Uganda chimeric virus in the presence or absence of trypsin and a blocking antibody against human DPP4. A) Fluorescent microscopy showing RFP expression 24 hours post-infection for each treatment group. B) Western blot of N protein and actin 24 hours post-infection.

### MERS-CoV therapeutics are ineffective against MERS-Uganda spike

Having established replication capacity in human cells, we next sought to determine if therapeutics developed against the MERS-CoV spike could disrupt infection with the MERS-Uganda spike chimera. Several monoclonal antibodies have been identified as possible therapeutic options for treatment of MERS-CoV, including LCA60 and G4. We first evaluated LCA60, a potent antibody that binds adjacent to the spike RBD of MERS-CoV (23). However, the major changes in the RBD region of MERS-Uganda spike predicted a lack of efficacy (**Fig. 4A**). LCA60 potently neutralized wild-type MERS-CoV grown in both the presence and the absence of tyrpsin (**Fig. 4B**). However, consistent with expectations, the LCA60 antibody had no impact on infection with the MERS-Uganda chimera, failing to neutralize the bat spike-expressing virus (**Fig. 4B**). We subsequently examined a second monoclonal antibody, G4, which had previously mapped to a conserved portion of the S2 region of the MERS-spike (**Fig. 4A**) (24). With the epitope relatively conserved in MERS-Uganda spike, we tested the efficacy against the zoonotic spike chimera. However, the results demonstrate no neutralization of MERS-Uganda spike virus by the S2-targeted antibody (**Fig. 4C**). Notably, G4 also failed to neutralize wild-type MERS-CoV grown in the the presence of exogenous trypsin (**Fig. 4C**). Together, the results indicate that both group 2C CoV spikes could escape neutralization by the S2-targeted antibody in the presence of exogenous trypsin. Overall, these experiments suggest that antibodies targeted against MERS-CoV, even to regions in the highly conserved S2 domain, may not have utility against viruses expressing the PDF-2180 spike.

**Figure 4.**
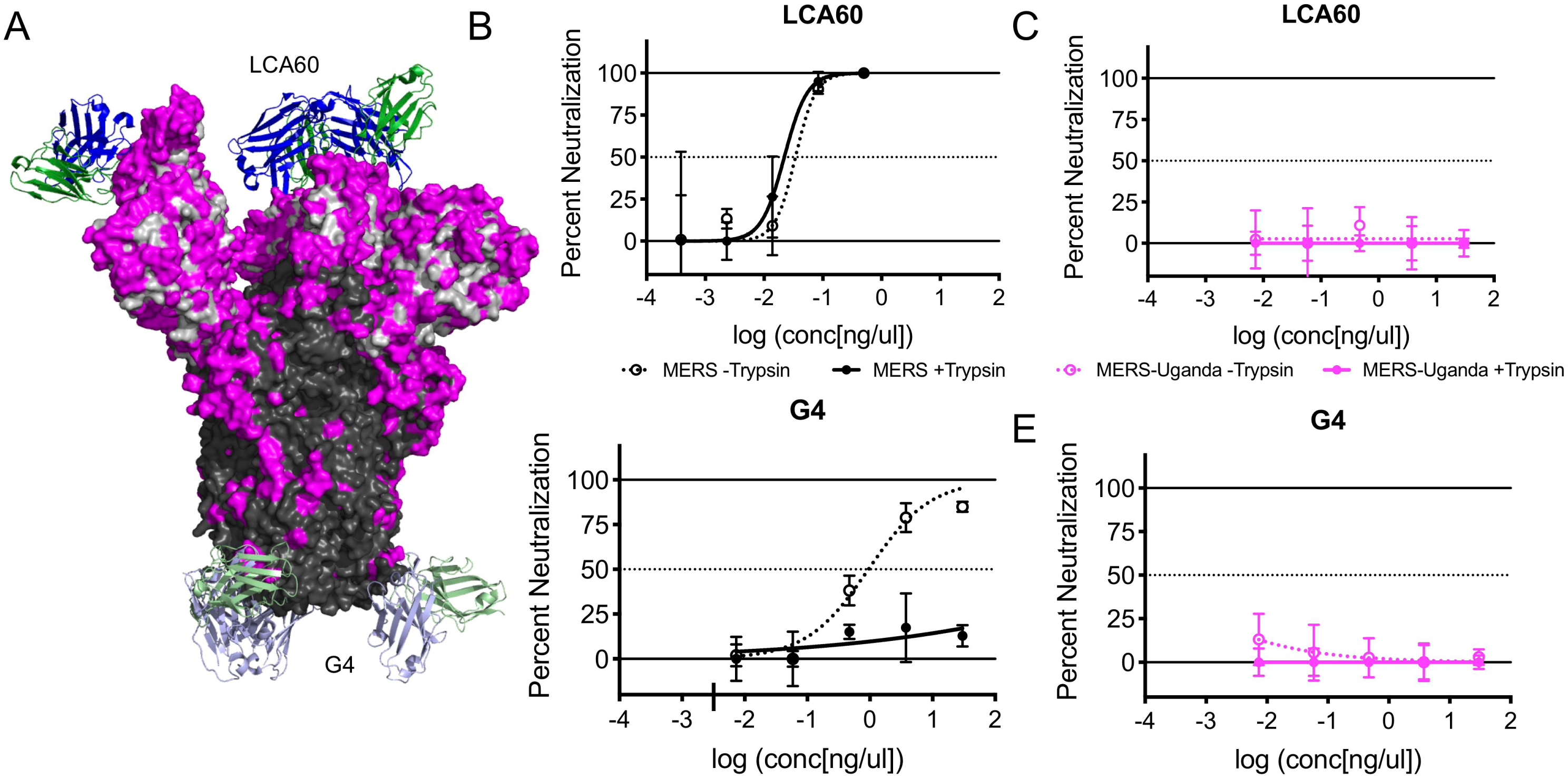
Antibodies against MERS-CoV fail to neutralize MERS-Uganda chimera. A) Structure of the MERS-CoV spike trimer with therapeutic antibody LCA60 bound adjacent to the receptor-binding domain and the antibody G4 bound to the S2 portion. Consensus amino acids are outlined for the S1 (grey) and S2 (black) domains, with PDF-2180 differences noted in magenta. B & C) Plaque neutralization curves for B) LCA60 and C) G4 with (solid) and without (dotted) trypsin treatment for MERS-CoV (black) and MERS-Uganda chimera (magenta) (n=3 per concentration).

### Trypsin treatment rescues the replication of zoonotic HKU5-CoV

Based on the MERS-Uganda chimera virus, we wondered if a similar barrier prevented replication of other zoonotic CoVs. Previously, our group had generated a full-length infectious clone for HKU5-CoV, another group 2C coronavirus sequence isolated from bats. Similar to the MERS-Uganda chimera, the infectious clone of HKU5-CoV produced sub-genomic transcripts, but failed to achieve productive infection (6). Revisiting the full-length recombinant virus, we sought to determine if trypsin treatment could also rescue HKU5-CoV. Following HKU5-CoV infection, addition of trypsin to the media resulted in cytopathic effect and cell fusion. In contrast, cultures lacking trypsin showed no signs of viral infection. Exploring viral genomic RNA, trypsin in the culture media permitted robust infection with HKU5-CoV that increased over time and was absent in cells not treated with trypsin (**Fig. 5A**). Similarly, trypsin in the media also permitted the accumulation and proteolytic cleavage of the HKU5 spike protein in a dose and time dependent manner (**Fig. 5B**). Importantly, the addition of anti-DPP4 antibody had no impact on HKU5-CoV infection, suggesting the use of a different receptor than used by wild-type MERS-CoV, similar to the findings with MERS-Uganda spike (**Fig. 5C**). Together, these results demonstrate that protease cleavage is a primary barrier to infection of Vero cells with HKU5-CoV.

**Figure 5.**
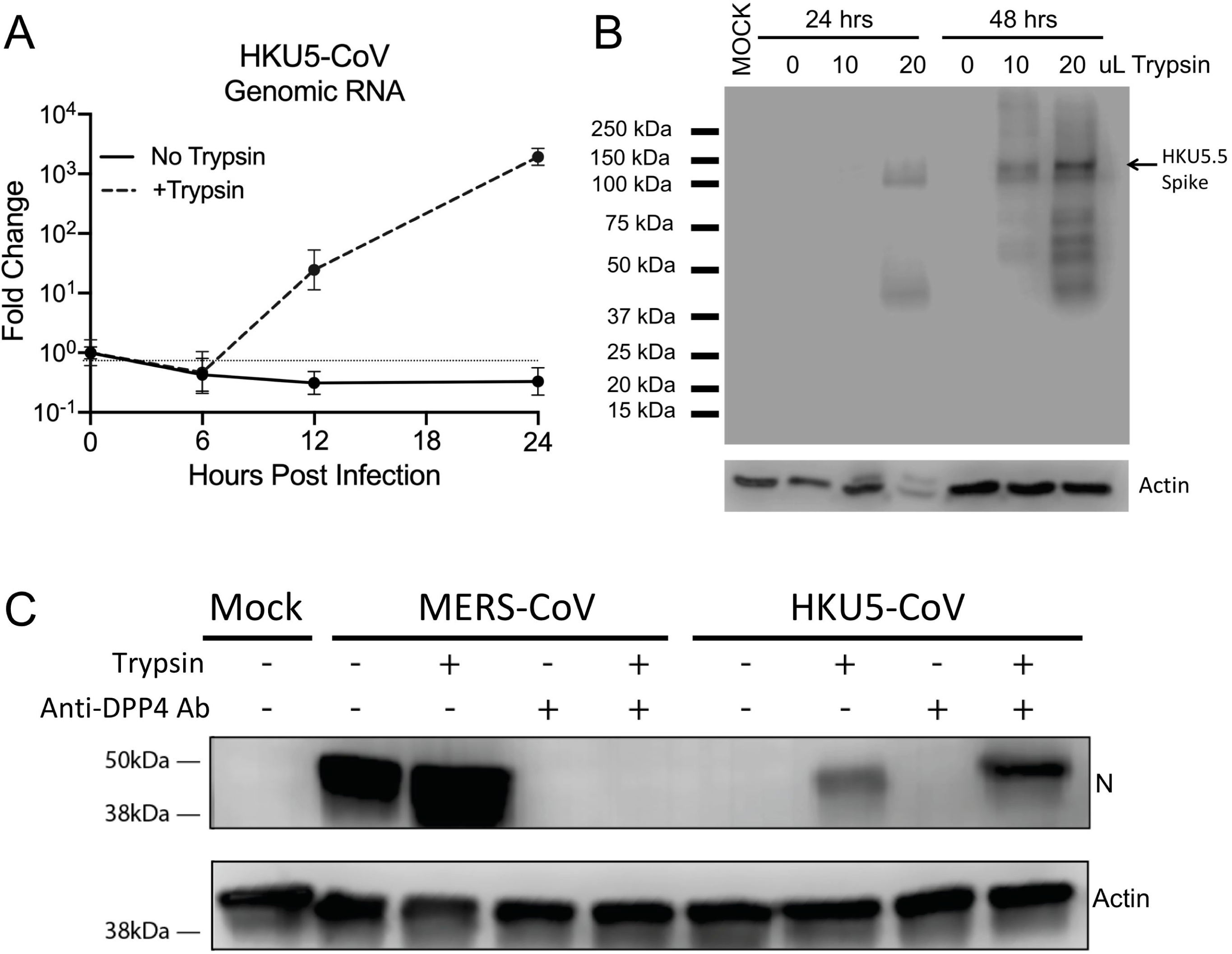
Exogenous trypsin rescues replication of HKU5-CoV. Vero cells were infected with full-length HKU5-CoV in the presence or absence of trypsin. A) Expression (qRT-PCR) of HKU5-CoV viral genome in the presence or absence of trypsin (n=3). B) Immunoblotting of HKU5 spike protein and cellular actin 24 and 48 hours post-infection with varying concentrations of trypsin in the media. C) Immunoblotting for MERS N protein and cellular actin following infection in the presence or absence of trypsin and human DPP4 antibody.

## Discussion

In this manuscript, we expanded our examination of circulating zoonotic viruses and identified protease cleavage as an important barrier to emergence of some group 2C zoonotic CoVs. The chimeric virus containing the spike protein from PDF-2180 was capable of replication in Vero cells and human cells (Huh7, Caco-2) if treated with exogenous trypsin. However, neither continuous nor primary human airway cultures were susceptible to infection, contrasting wild-type MERS-CoV. The MERS-Uganda chimera also maintained replication despite treatment with antibodies blocking human DPP4, suggesting use of either an alternative receptor or a different entry mechanism for infection. Importantly, current therapeutics targeting the MERS spike protein showed no efficacy against the MERS-Uganda chimera, highlighting a potential public health vulnerability to this and related group 2C CoVs. Finally, the trypsin-mediated rescue of a second zoonotic group 2C CoV, HKU5-CoV, validates findings that suggested that protease cleavage may represent a critical barrier to zoonotic CoV infection in new hosts (25, 26). Together, the results highlight the importance of spike processing in CoV infection, expand the correlates associated with emergence beyond receptor binding alone, and provide a platform strategy to recover previously non-cultivatable zoonotic CoVs.

With the ongoing threat posed by circulating zoonotic viruses, understanding the barriers for viral emergence represents a critical area of research. For CoVs, receptor binding has been believed to be the primary constraint to infection in new host populations. Following the SARS-CoV outbreak, emergence in humans was attributed to mutations within the receptor-binding domain that distinguished the epidemic strain from progenitor viruses harbored in bats and civets (27). Yet, work by our group and others has indicated that zoonotic SARS-like viruses circulating in Southeast Asian bats are capable of infecting human cells by binding to the known human ACE2 receptor without adaptation (13, 14, 28). Similarly, pseudotyped virus studies have identified zoonotic strains HKU4-CoV and NL140422-CoV as capable of binding to human DPP4 without mutations to the spike (26, 29). In this study, we demonstrate that both PDF-2180 and HKU5-CoV spikes are capable of binding to and infecting human cells if primed by trypsin cleavage. Together, the results argue that several circulating zoonotic CoV strains have the capacity to bind to human cells without adaption and that receptor binding may not be the only barrier to CoV emergence.

Data from this study implicates the processing of the spike protein as a critical factor for CoV infection. In the absence of trypsin, the MERS-Uganda and HKU5-CoV spikes were unable to mediate infection and initially suggested a lack of receptor compatibility (6, 16). However, exogenous trypsin treatment produced robust infection, indicating that despite binding to human cells, CoVs cannot overcome incomplete spike processing. As such, evaluating zoonotic virus populations for emergence threats must also consider the capacity for CoV spike activation in addition to receptor binding. In this new paradigm, the combination of receptor binding and proteolytic activation by endogenous proteases permits zoonotic CoV infection, as with MERS-CoV and SARS-CoV (**Fig. 6**). The absence of receptor binding (**Fig. 6A**) or compatible host protease activity (**Fig. 6B**) restricts infection with certain zoonotic strains like PDF-2180 or HKU5-CoV. These barriers can be overcome with the addition of exogenous proteases, disrupting the need for host proteases and permitting receptor-dependent or receptor-independent entry (**Fig. 6C**). Overall, the new paradigm argues that both receptor binding and protease activation barriers must be overcome for successful zoonotic CoV infection of a new host.

**Figure 6.**
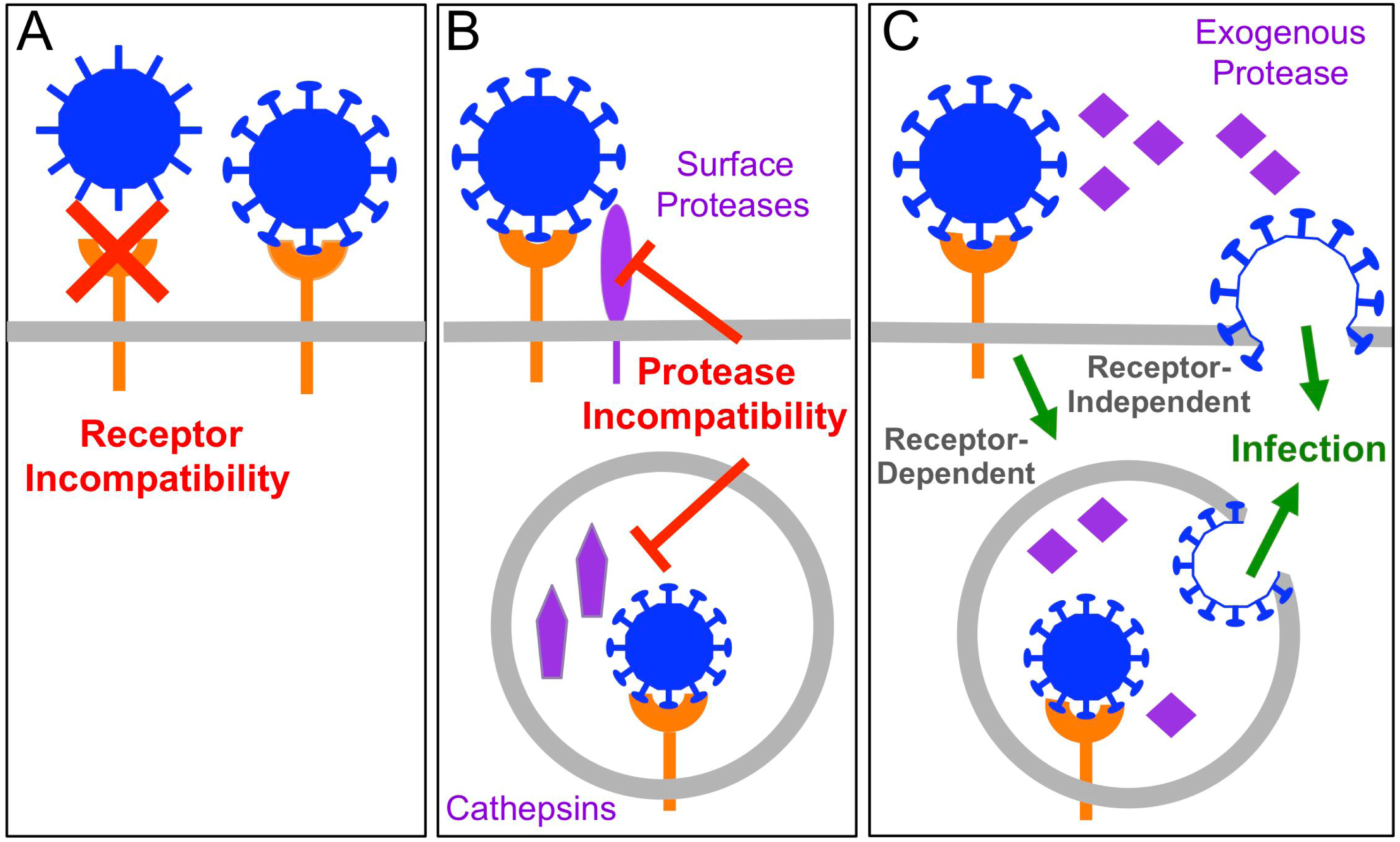
Barriers to zoonotic coronavirus emergence. Both receptor binding and protease activitation are key correlates that govern zoonotic coronavirus emergence. A) A lack of receptor binding with zoonotic CoVs precludes the infection of new host cells. B) Despite receptor binding, the absence of compatible host proteases for spike cleavage restricts infection in new hosts. C) The addition of exogenous protease overcomes the host protease barriers and may or may not require receptor binding.

The requirement for exogenous trypsin treatment is not unique to MERS-Uganda or HKU5-CoV. Influenza strains are well known to require trypsin treatment to facilitate their release in cell culture (30). In addition, highly pathogenic avian influenza strains have been linked to mutations that improve cleavage by ubiquitous host protese, augmenting their tissue tropism and virulence (31). Similarly, a wealth of enteric viruses, including polio, cowpox, and rotaviruses, depend on trypsin to prime, modulate, and/or expand infection (32, 33). Even within the CoV family, enteric viruses, including PEDV, porcine delta CoV, and swine acute diarrhea syndrome (SADS) CoV require trypsin for replication *in vitro* (34-36). Together, these prior studies illustrate the importance of protease activation in virus infections. However, the protease barrier to PDF-2180 and HKU5-CoV spike-mediated infection may also reflect on the emergence of SARS-CoV and MERS-CoV. While initial studies argued that receptor binding was the primary barrier, the existence of zoonotic strains capable of efficiently using the same human entry receptors contradicts that suggestion (13, 14). It is possible that emergence of epidemic CoV strains also requires modifying protease cleavage in either humans or an intermediate host, such as camels or civets, in addition to increased receptor-binding affinity. Consistent with this idea, reports have detailed differential infection with MERS-CoV based on host protease expression (18). Similarly, mouse adaptation of MERS-CoV resulted in spike modifications that alter protease activation and entry *in vivo* (37). While group 2B bat CoV strains (WIV1-CoV, WIV16-CoV and SHC014-CoV) do not require trypsin for infection (9, 13, 14, 38), differences in protease activation may contribute to infection changes relative to the epidemic SARS-CoV. In this context, our findings expand the importance of protease cleavage as a criterion to consider for zoonotic virus emergence in a new host population.

In evaluating the threat to humans posed by PDF-2180 and HKU5-CoV, the results demonstrate a pathway to emergence. Neither CoV spike uses human DPP4 for entry, and the PDF-2180 chimera failed to replicate in human respiratory models, even in the presence of trypsin. However, replication in Huh7 and Caco-2 cells indicates human infection compatibility and may portend differential tropism, possibly in the alimentary or biliary tracts, as has been described for several mammalian CoVs (34-36). MERS-Uganda or HKU5-CoV could utilize this same trypsin-rich environment in the gut to emerge as an enteric pathogen in humans, although its pathology and virulence would be hard to predict. Evidence from both SARS-CoV and MERS-CoV outbreaks suggests the involvement of enteric pathways during infection (39, 40). Replication in the gut might select for mutations that expand spike processing/tropism and allow replication in other tissues, including the lung, and lead to virulent disease in the new host population, as seen with Porcine Respiratory Coronavirus (41). In examining the threat posed by PDF-2180 and HKU5-CoV, we must consider the emergence of these CoVs in tissues other than the lung and harboring distinct pathologies compared to epidemic SARS and MERS-CoV.

The receptor dynamics of MERS-Uganda and HKU5-CoV also remain unclear in the context of this study. In the presence of trypsin, neither spike protein requires the MERS-CoV receptor, DPP4 for entry, which is consistent with the differences between the receptor-binding domains of the bat and epidemic strains. Therefore, it was not surprising that antibodies that target the RBD of the MERS-CoV spike were ineffective in blocking infection of the PDF-2180 chimera. However, the S2-targeted antibody, G4, also had no efficacy against MERS-Uganda, despite a relatively conserved binding epitope. This result is possibly explained by differing amino acid sequences between MERS-CoV and PDF-2180 at the G4 epitope, specifically residue 1175, which is associated with G4 escape mutants in MERS-CoV (24). Alternatively, the G4 antibody also failed to neutralize wild-type MERS-CoV grown in the presence of trypsin, indicating that entry is still possible, despite treatment with antibody binding the S2 domain. Conversely, the presence of trypsin may prime a receptor-independent entry for the MERS-Uganda chimera, similar to the JHVM strain of MHV (42). Yet, this result would contrast PEDV, which requires receptor binding prior to trypsin activation to facilitate infection (35). Importantly, the lack of infection in respiratory cells suggests that some receptor or attachment factor is necessary to mediate entry with the PDF-2180 spike. Recent work with MERS-CoV binding sialic acid supports this idea (43) and indicates that the PDF-2180 spike may not have a similar binding motif. Ovrall, further experimental studies are required to fully understand the receptor dynamics of the PDF-2180 spike.

While providing a new strategy to recover zoonotic CoVs, this manuscript highlights proteolytic cleavage of the spike as a major barrier to group 2C zoonotic CoV infection. For both MERS-Uganda and HKU5-CoV, the addition of exogenous trypsin rescues infection, indicating that spike cleavage, not receptor binding, limits these strains in new hosts and tissues. The adaptation of the protease cleavage sites or infection of tissues with robust host protease expression could permit these two zoonotic CoV strains to emerge and may pose a threat to public health due to the absence of effective spike-based therapeutics. In considering cross-species transmission, our results using reconstructed bat group 2C CoVs confirm spike processing as a correlate associated with emergence. Adding spike processing to receptor binding as primary barriers offers a new framework to evaluate the threat of emergence for zoonotic CoV strains.

## Methods

### Cells, viruses, in vitro infection, and plaque assays

Vero cells were grown in DMEM (Gibco, CA) supplemented with 5% FetalClone II (Hyclone, UT) and antibiotic/antimycotic (anti/anti) (Gibco, CA). Huh7 cells were grown in DMEM supplemented with 10% FetalClone II and anti/anti. Caco-2 cells were grown in MEM (Gibco, CA) supplemented with 20% Fetal Bovine Serum (Hyclone, UT) and anti/anti. Human airway epithelial cell (HAE) cultures were obtained from the UNC CF Center Tissue Procurement and Cell Culture Core from human lungs procured under Univeristy of North Carolina at Chapel Hill Institutional Review Board-approved protocols. Wild-type MERS-CoV, chimeric MERS-Uganda and HKU5-CoV were cultured on Vero cells in OptiMEM (Gibco, CA) supplemented with anti/anti. For indicated experiments, trypsin (Gibco, CA) was added at 0.5 µg/ml unless otherwise indicated.

Generation of wild-type MERS-CoV, MERS-Uganda, and HKU5-CoV viruses utilized reverse genetics and have been previously described (6, 17, 44). For MERS-Uganda chimera expressing RFP, we utilized the MERS-CoV backbone, replacing ORF5 with RFP as previously described (17). Synthetic constructions of chimeric mutant and full-length MERS-Uganda and HKU5-CoV were approved by the University of North Carolina Institutional Biosafety Committee. Replication in Vero, Calu-3 2B4, Caco-2, Huh7, and HAE cells was performed as previously described (12, 45-47). Briefly, cells were washed with PBS and inoculated with virus or mock diluted in OptiMEM for 60 minutes at 37 °C. Following inoculation, cells were washed 3 times, and fresh media with or without trypsin was added to signify time 0. Three or more biological replicates were harvested at each described time point. For HAE cultures, apical surfaces were washed with PBS containing 5ug/ml trypsin at 0, 8, 18, 24, and 48 hours post infection. No blinding was used in any sample collections, nor were samples randomized. Microscopy photos were captured via a Keyence BZ-X700 microscope.

For antibody neutralization assays, MERS-CoV and MERS-Uganda stocks were grown in OptiMEM both with and without trypsin. All stocks were quantified via plaque assay by overlaying cells with 0.8% agarose in OptiMEM supplemented with 0.5 µg/ml trypsin and anti/anti. MERS-Uganda stocks grown without trypsin had low titers but were sufficient for neutralization assays.

For anti-DPP4 blocking experiments, Vero cells were preincubated with serum-free OptiMEM containing 5ug/ml anti-human DPP4 antibody (R & D systems, MN) for one hour. Media was removed and cells were infected for 1 hour with virus or mock inoculum at a multiplicity of infection of 0.1. The inoculum was removed, cells were washed three times with PBS, and media was replaced.

### RNA isolation and quantification

RNA was isolated via TRIzol reagent (Invitrogen, CA) and Direct-zol RNA MiniPrep kit (Zymo Research, CA) according to the manufacturer’s protocol. MERS-CoV and MERS-Uganda gRNA was quantified via TaqMan Fast Virus 1-Step Master Mix (Applied Biosystems, CA) using previously reported primers and probes targeting ORF1ab (47) and normalized to host 18S rRNA (Applied Biosystems, CA). HKU5-CoV RNA was first reverse transcribed using SuperScript III (Invitrogen) and was, CA) then assayed using SsoFast EvaGreen Supermix (Bio-Rad, CA) and scaled to host GAPDH transcript levels. HKU5 gRNA was amplified with the following primers: Forward – 5’-CTCTCTCTCGTTCTCTTGCAGAAC-3’, Reverse – 5’-GTTGAGCTCTGCTCTATACTTGCC-3’. GAPDH RNA was amplified with the following primers: Forward – 5’-AGCCACATCGCTGAGACA- −3’, Reverse – 5’-GCCCAATACGACCAAATCC-3’. Fold change was calculated using the ΔΔCt method and was scaled to RNA present at 0 hours post-infection.

### Generation of VRP, polyclonal mouse antisera, and western blot analysis

Virus replicon particles (VRPs) expressing the MERS-CoV nucleocapsid or HKU5-5 CoV spike were constructed using a non-select BLS2 Venezuelan Equine Encephalitis (VEE) virus strain 3546 replicon system as previously described (48). Briefly, RNA containing the nonstructural genes of VEE and either MERS-CoV nucleocapsid or HKU5-5 CoV spike was packaged using helper RNAs encoding VEE structural proteins as described previously (49). Six-week-old female BALB/c mice were primed and boosted with VRPs to generate mouse anti-sera towards either MERS-CoV nucleocapsid or HKU5-5 CoV spike. Following vaccination, mouse polyclonal sera were collected as described previously (50). For Western blotting, lysates from infected cells were prepared as described before in detail (51), and these blots were probed using the indicated mouse polyclonal sera. MERS-CoV N sera was able to detect to HKU5-CoV N protein via Western blot as previously described (7).

### Virus neutralization assays

Plaque reduction neutralization titer assays were preformed with previously characterized antibodies against MERS-CoV as previously described (23, 24). Briefly, antibodies were serially diluted 6- to 8-fold and incubated with 80 PFU of the indicated viruses for 1 h at 37°C. The virus and antibodies were then added to a 6-well plate of confluent Vero cells in triplicate. After a 1 hour incubation at 37°C, cells were overlaid with 3 ml of 0.8% agarose in OptiMEM supplemented with 0.5 µg/ml trypsin and anti/anti. Plates were incubated for 2 or 3 days at 37°C for MERS-CoV or MERS-Uganda, respectively, and were then stained with neutral red for 3 h, and plaques were counted. The percentage of plaque reduction was calculated as [1 - (no. of plaques with antibody/no. of plaques without antibody)] × 100.

### Biosafety and biosecurity

Reported studies were initiated after the University of North Carolina Institutional Biosafety Committee approved the experimental protocols. All work for these studies was performed with approved standard operating procedures (SOPs) and safety conditions for MERS-CoV and other related CoVs. Our institutional CoV BSL3 facilities have been designed to conform to the safety requirements recommended in Biosafety in Microbiological and Biomedical Laboratories (BMBL), the U.S. Department of Health and Human Services, the Public Health Service, the Centers for Disease Control (CDC) and the National Institutes of Health (NIH). Laboratory safety plans have been submitted, and the facility has been approved for use by the UNC Department of Environmental Health and Safety (EHS) and the CDC.

## Acknowledgments

The research described in this manuscript was supported by grants from the United States Agency for International Development (USAID) Emerging Pandemic Threats PREDICT project (cooperative agreement number GHN-A-OO-09-00010-00) and from the National Institute of Allergy & Infectious Disease and the National Institute of Aging of the NIH under awards U19AI109761 and AI110700 to RSB; R00AG049092 to VDM. HAE cultures were supported by the National Institute of Diabetes and Digestive and Kidney Disease under award NIH DK065988 to SHR. Trypsin resistant Vero cells were kindly provided by Linda Saif. Monoclonal antibody LCA60 was provided by Davide Corti and Humabs Biomed SA. The content described herein is solely the responsibility of the authors and does not necessarily represent the official views of the NIH.

